# Health-seeking behavior and associated factors among community in Southern Ethiopia:Community based cross-sectional study guided by Health belief model

**DOI:** 10.1101/388769

**Authors:** Likawunt Samuel Asfaw, Samuel Yohannes Ayanto, Yitagessu Habtu Aweke

## Abstract

**Background:** Health-seeking behavior is a sequence of actions taken to promote health and prevent disease. Governments’ expenditure to health is being improved in Ethiopia. In contrast, high disease burden and low health service utilization is observed. The low health service utilization could be due to low health-seeking behavior of the community. Thus, this studywas aimed to determine the level of health-seeking behavior and associated factors in Hosanna town, Southern Ethiopia.

**Methods:** We used communitybased cross-sectional study design among community (n= 443) in Hosanna town. The overall health-seeking behavior of study participants was assessed using the mean score of each of the dimensions (health promotion and disease prevention activities) as a cut-off value. Having a score above the mean on each of the target dimensions was equated with having a high level of health seeking behaviour. STATA 12 soft-ware package (Stata Corporation, College Station, Texas, 77845, USA)was used for descriptive and logistic regression analysis.

**Results:** About eighty five percent of (85.4%) participants had low level of health-seeking behavior. Males were about two folds more likely to have low level of health-seeking behavior than females (AOR: 1.8; CI 1.03–3.42). Widowed participants were about five times more likely to have low health-seeking behavior (AOR: 4.8; CI 2.1–17.1) when compared to married participants. Those participants who are illiterate were about five times more likely to have low health-seeking behavior than who completed higher education (AOR: 4.5; CI 1.16–17.8).

**Conclusion:** The study revealed low health-seeking behavior among the study participants in the study area. This finding suggested the need forurgent interventions to the health literacy packages of Ethiopia to enhance the health seeking behavior of the country.

## Introduction

Health is a comprehensive concept that encompasses all social and biological aspects of life and health-seeking behavior refers to a sequence of actions taken to promote health and prevent disease [1]. The health policies and strategies entail knowledge about health seeking behavior for health promotion, disease prevention and improve quality of life [1, 2]. The practice of health seeking behaviour has a marvelous potential to reduce the occurrence of disease, disability and death [1]. Healthcare utilization that is an immediate outcome of health seeking behaviour is also found to be important to get health counseling (family planning, antenatal care, growth monitoring), screening for chronic diseases, and adherence to effective treatment [3–5]. However, we have learnt the growing of evidences in low health care utilization and increased disease burden [6–10].

Previous studies showed that Africa accounts for 22% of global disease burden and adult mortality rate 347 per 1000 population, which is the highest in the world [4]. Similarly, despite an improved access in physical health facilities and government expenditure on health, the burden of diseases was not successfully reduced and health service coverage is low in Ethiopia [7–11]. For instance, according to the WHO African health observatory report, the burden of diseases was 73.6% in Ethiopia in 2010 exceeding the African average (71.1%) [7]. Moreover, 350 premature deaths from all causes and 14.8 crude death rate per 1000 of population was recorded in 2010 which is the highest in east Africa [9]. Concerning maternal health, one in 26 African mothers have chances of dying due to preventable pregnancy related complications. This is about 300 times higher than reported for developed nations [4].

Regarding service utilization, the Ethiopian demographic and health survey finding of 2011 reported low level of health service utilization. More than four in five mothers did not receive antenatal care and only 10% of ill individuals due to all cause got treatment [8–10]. To the best of our understanding, the observed evidences does not reflect nature more deserving for high income countries than low income countries, rather the difference in level of health seeking behaviour in high and low income countries.

Ethiopia is currently experiencing an incidence of newly emerging and reemerging health problems and in state of transition [2, 3], that requires comprehensive health care policies and programs. Similarly, unlike to the traditional assumptions chronic non communicable (Diabetes Mellitus, Cardiovascular disorders, cancer) illnesses are not confined to people of developed countries [12–15].

Evidences show that socio-economic status, geographic settings, cultural issues, service quality, health system policy and procedures are among the factors affecting health-seeking behavior of the community [15–18]. Individuals who fail to get health information found to have lower health seeking behavior [17]. However, individuals having higher health seeking behavior could better prevent disease and promote health [18–20].Healthcare-seeking behavior is a multifaceted effect and needs an appropriate investigation in order to provide knowledge that will help with the formulation of health care policies and programs. Thus, in this study we assessed the level of health seeking behaviors and associated factors in urban households.

## Methods

### Study area and period

The study was conducted in Hosanna town, the capital of Hadiya Zone, located at a distance of 232 kilometers southwest of Addis Ababa, the capital of Ethiopia. There were 16,707 Households in the town [20].Community based cross-sectional study was carried out among residents of the town, in August 2013

### Sample size and sampling technique

The sample size for our study was estimated by taking prevalence of health seeking behaviour (50%), 95% confidence level, 5% margin of error and 15% non-response rate. Consequently, the final sample size was determined to be 443participants.

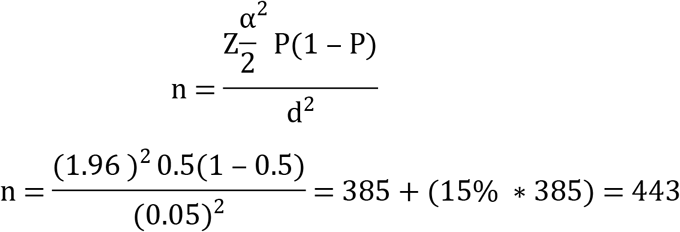

The final sample size was proportionally allocated to sub-administrative units of the town. Sampling frame was created for each sub-unit and randomly generated numbers were used to select the households. Simple random sampling technique was used to selectthe households from each unit. From each of the selected households the head of the house hold was included in data collection.

### Data collection instrument and procedures

The health seeking questionnaire, WHO STEPS instrument and global physical activity questionnaire (GPAQ) were modified and used for data collection. The modified instrument was translated in to the local language, Amharic. Data were collected through self-administered using structured questionnaire. Following informed oral consent procedures, the head of the households were asked to complete at home level in a quiet corner away from the presence of other people and drop to box prepared. Each took approximately fifty minutes. Probes and clarifications were sought as deemed necessary. Two days training was provided for data collectors and supervisors regarding research ethics, data collection procedures and contents of the instrument to increase the quality of our data. Supportive supervision was carried out by the supervisors on a daily basis during the data collection period. The completed questionnaire had been checked daily for its completeness and consistency.

### Variables and its measurement

The dependent variable was health seeking behavior. It is a composite variable measured using model construct. The overall health-seeking behavior of study participants was assessed using the mean score of each of the following four dimensions;

#### Actions taken when got ill

The health seeking behaviour of participants for this dimension was obtained from the following questions; ‘during your last illness did you seek treatment?’ This question had the following response categories: ‘Yes if seek care from health facilities and or traditional healers’, ‘No did not seek treatment’.

#### Screening for general health

This dimension was assessed using the following questions with “Yes -coded as 1” or “No -coded as 0” responses; ‘have you ever checked your blood pressure to know the level of your blood pressure?, ‘Did you ever checked your blood sugar level to know the level of your blood sugar?’, ‘Have you ever tested for human immune deficiency virus (HIV) infection for early care and treatment?,’ ‘ Did you vaccinated children and any family member who is eligible for?’, ‘ Did you or member of your family monitor the growth of recent child in family?’, Did you or member of your family followed antenatal care for the resent pregnancy?’ In this dimension participants having a score above the mean was equated with having high health seeking behaviour of screening for general health were coded as “Yes 1” and otherwise coded as ‘No 0’, indicating that they had low health seeking behaviour of screening for general health.

#### Health oriented leisure activities

The health seeking behaviour of participants of health oriented leisure activities was measured as high ‘Yes 1’ -if scored above the mean for questions of aerobic physical activities (walking, running, swimming, and bicycling) and health oriented leisure activities (playing tennis, jumping rope, lifting weight) or low level of health seeking for health oriented leisure activities ‘No 0’-if scores below the mean.

#### Risk exposure

Health seeking behaviour of participants of risk exposure disease was measured using the question, “Did you take alcohol?”, “Did you smoke tobacco products?”, and “Did you chew Khat?” with “yes” or “no” responses in both. Participants responded “Yes” to at least one of these questions were coded as ‘0’, indicating that they had high risk exposure and low health seeking behaviour.

Overall, having a score above the mean on each of the target dimensions was equated with having a high level of health seeking behaviour. The exposure variables included age, sex, education, occupation, marital status, family income, and distance from health facility.

### Data analysis techniques

The collected data were cleaned and entered to Epi-Data version 3.2, and exported to STATA 12 soft-ware package (Stata Corporation, College Station, Texas, 77845, USA)for analysis. Descriptive statistics and multivariable logistic regression were used to analyze the data. Candidate variables with P-value <0.2 in Bivariable model were entered to multivariable model to adjust for predictors. The 95% CI for the corresponding Odds Ratio (OR) was used to assess the degree of associations at (P<0.05) to declare significance.

### Definition of terms

#### High health seeking behaviour

Participants having a score above (≥) the mean on each of the target dimensions was equated with having a high level of health seeking behaviour

#### Low health seeking behaviour

Participants having a score below the mean on each of the target dimensions was equated with having a high level of health seeking behaviour

#### Ethics

Institutional research ethics review committee of Hosanna College of Health Sciences approved and granted permission of this study. Informed verbal consent was obtained from all study participants before data collection after explaining the objectives of the research. In this research we obtained informed verbal consent from the research participants because all the data sought was associated purely with information rather than human samples or did not put participants on experiment, which needs national ethical approval in our context. We obtained ethical clearance for the research to be conducted in this way. This is the reason why we obtained informed verbal consent than written.

## Results

### Characteristics of study participants

Total of 443 questionnaires were received, only 424 questionnaires were valid and included in analysis. Of the questionnaires deemed not valid, 4 respondents refused and the rest did not complete the questionnaire in that one subsection of the questionnaire omitted. The mean age of the study participants was 33.8 ± 11.4 Standard deviation (SD) years. Pertaining to family size, in average five people living in one house. Of the 424 participants 51.3 % were males and 61.6% were currently married [table 1–2].

**Table 1:** Socio-demographic characteristics of study participants

| Variables | Frequency | Percent (%) |
| --- | --- | --- |
| **Sex** |  |  |
| Male | 218 | 51.4 |
| Female | 206 | 48.6 |
| **Marital status** |  |  |
| Married | 262 | 61.8 |
| Single | 84 | 19.8 |
| Divorced | 45 | 10.6 |
| Widowed | 33 | 7.8 |
| **Level of education** |  |  |
| No formal education | 55 | 13 |
| Primary education | 162 | 38.2 |
| Secondary education | 109 | 25.7 |
| Higher education | 98 | 23.1 |
| Total | 424 | 100.00 |

**Table 2:** Socio-demographic characteristics of study participants

| Characteristics | Frequency | Percent (%) |
| --- | --- | --- |
| **Occupation** |  |  |
| Government employee | 144 | 34.0 |
| Non-Government employee | 75 | 17.7 |
| Merchant | 117 | 27.6 |
| Farmer | 23 | 5.4 |
| Others | 65 | 15.3 |
| **House family live in** |  |  |
| Own | 282 | 30.4 |
| Rent | 129 | 66.5 |
| Other | 13 | 3.1 |
| **Monthly income** |  |  |
| <18.5 USD | 75 | 17.7 |
| 18.5-36.5 USD | 177 | 41.7 |
| 36.6-72.5 USD | 120 | 28.3 |
| >72.6 USD | 52 | 12.3 |
| Total | 424 | 100 |

### Health seeking behaviors

Table 3 shows health seeking behaviour of study participants. Accordingly, 397 (93.6%) took action when got ill. Of those who took action when got ill, 345 (86.9%) visited medical institutions, and 24(6.0%) use only traditional home remedies. Of the participants visited medical institutions, 139 (40.2%) preferred private clinics as their first choice, 106(30.4%) sought health care from Hospital, 82(23.7%) sought health care from pharmacy and only 18 (5.4%) sought health care from public health centers.

The study participants were also asked to rate their perceived health status and the self-rated perceived health status show 113 (26.7%) of participants felt very well, 110(25.9%) felt excellent and 17(4.0%) were unable to express their health status.

Only 72 (17.0%) of participants undertook screening for general health status and 352 (83.0%) of study participants did not to screening. Similarly, more than eighty percent (81.6%) participants did not undertook health oriented leisure activities. The general prevalence of health seeking behavior was 14.6% and 85.4% of study participants had low level of health seeking behaviour [table 4].

**Table 3:** Health-seeking behavior of study participants in Hosanna

| Characteristics | Number | Percent (%) |
| --- | --- | --- |
| **Perceived health status of participants** |  |  |
| Excellent | 110 | 25.6 |
| Very good | 113 | 26.7 |
| Good | 100 | 23.6 |
| Fair | 51 | 12.0 |
| Bad | 33 | 7.8 |
| Difficult to express | 17 | 4.0 |
| **Take action when got ill** |  |  |
| Yes | 397 | 93.6 |
| No | 27 | 6.4 |
| **Reported actions taken when got ill** |  |  |
| Visiting medical institutions | 345 | 86.9 |
| Use traditional medicine and home remedies | 24 | 6.0 |
| Combination of both | 28 | 7.1 |
| **Medical institution of first choice** |  |  |
| Private clinics | 139 | 40.2 |
| Hospital | 106 | 30.7 |
| Health centers | 18 | 5.4 |
| Pharmacy | 82 | 23.7 |
| **Total** | **424** | **100.0** |

**Table 4:** Health-seeking behavior of study participants in Hosanna

| Characteristics | Number | Percent (%) |
| --- | --- | --- |
| **Take health oriented leisure activities** |  |  |
| Yes | 78 | 18.4 |
| No | 346 | 81.6 |
| **Reported reasons not to take leisure** |  |  |
| Lack of knowledge | 157 | 45.4 |
| Lack of time | 81 | 23.4 |
| Lack of access | 87 | 25.1 |
| Others | 21 | 6.1 |
| **Screened for general health** |  |  |
| Yes | 72 | 17.0 |
| No | 352 | 83.0 |
| **Vaccination** |  |  |
| Yes | 393 | 92.7 |
| No | 31 | 7.3 |
| **Antenatal care for mother** |  |  |
| Yes | 401 | 94.6 |
| No | 23 | 5.4 |
| **Health-seeking behavior** |  |  |
| Low | 362 | 85.4 |
| High | 62 | 14.6 |
| **Total** | **424** | **100.0** |

### Factors associated with health seeking behaviors

Table 5 presents the logistic regression model fitted to assess factors associated with health seeking behavior. Accordingly, marital status, sex and level of education were independently associated with health seeking behavior.

The odds of low health seeking behavior among widowed participants was 4.8 times higher than single ones (AOR = 4.8, CI: 2.1, 17.1). As shown in the adjusted model, the likelihood of low health seeking behaviour was significantly higher in male than female (AOR = 1.8, CI: 1.04, 3.42).

Similarly, in this study the association between level of education and health seeking behaviour was statistically significant (P<0.05), indicating that participants having no formal education had low heath seeking behaviour compared to those having higher education (AOR = 4.5, CI: 1.16, 17.8).

**Table 5:** Variables associated with health-seeking behavior

| Variables | Health-seeking behavior |  | Multivariate OR<br>(95%CI) | P-value |
| --- | --- | --- | --- | --- |
|  | Low (%) | High (%) |  |  |
| **Sex** |  |  |  |  |
| Male | 184(89.3) | 22(10.7) | 1.8(1.04,3.42) | 0.02<sup>†</sup> |
| Female | 178(79.8) | 40(18.3) | 1 |  |
| **Marrital status** |  |  |  |  |
| Married | 218(83.2) | 44(16.8) | 1 |  |
| Single<sup>‡</sup> | 69(82.1) | 15(17.9) | 2.1(0.9,12.5) | 0.21 |
| Divorced | 43(95.6) | 2(3.0) | 4.2(1.1,21.1) | 0.03<sup>†</sup> |
| Widowed | 32(97.0) | 1(3.0) | 4.8(2.1,17.1) | 0.02<sup>†</sup> |
| **Level of education** |  |  |  |  |
| Can't write or read | 47(85.5) | 8(14.5) | 4.5(1.16,17.8) | 0.02<sup>†</sup> |
| Primary | 139(85.8) | 23(14.2) | 1.4(0.50,4.20) | 0.48 |
| Secondary | 89(81.7) | 20(18.3) | 2.1(0.8,5.7) | 0.12 |
| Higher education | 87(88.8) | 11(11.2) | 1 |  |

‡=Not ever married †=P-value <0.05 1=Reference

## Discussion

Many evidences suggest that addressing health seeking behavior pave ways for appropriate utilization of health care services [19]. This study tried to measure health-seeking behavior in multidimensional approaches to improve specific health behavior change to prevent disease and promote health. Based on our measure, the study showed that majority (85.4%) of participants had low level of health-seeking behavior. The extent of health-seeking behavior of the current study was remarkably low when compared todifferent parts of the world [1,2].This finding was consistent with findings reported for mothers’ health care seeking behavior for child health illness in Dera district, North Shewa zone in Oromiya regional state of Ethiopia [16]. In any case, this finding implies that significant behavioral interventions are needed to improve health-seeking behavior so that increase health service utilization coverage in the community.

About 93.6% participants in our study took actions and seek medical help when got ill, which is more than reported in South Africa, 76.5%[3]. Our data also showed that 40.3% of participants primarily chose private clinics when they seek medical help. This finding is similar with a study reported in Ethiopia [3].

A decision made against maternal and child health care utilization was also used as an indicator for health seeking behavior. Accordingly, the result showed that it was 94.6% for maternal and 92.7% for child health care conditions. This is incomparably higher than findings of demographic and health survey(DHS)of 2011 in Ethiopia [20]. This could be due to the fact that our sample consisted of participants entirely from the urban setting unlike DHS that encompas both urban and rural regions throughout the country.

Socio-demographic characteristics of household heads were tested for association. The results ilustrated that sex, marrital statusand level of education showed an association with health-seeking behavior. Male had low score of health-seeking behavior compared to their counterparts. This finding was in line with study reported in Stockholm [22, 24]. Socially, women are more responsible for their family, often stay longer time in home and take time to identify theirhealth problems. Observations claimed that women are sensitive to their health. Conversely, men stay longer time out of door and busy in social matters representing their family in Ethiopia.

Pertaining to the influence of marital status to health seeking behaviour, divorced and widowed participants had low health-seeking behavior in our study. This finding was consistent with study reported for Jamaica in 2009 [19, 25]. Participants reported lower level of education had low health-seeking behavior. Most of the reports from Ethiopia and other countries supported this finding [25–27]. Consistencies in findings imply the influence of level of education on health seeking behaviour.

The present study has some relevant limitations that impede the power. One of the limitations of this study is related to the cross-sectional study design, in which the temporal relationships between the outcome and predictor variables cannot be established. Moreover, the sample was limited to single population which can limit the power of the study. We recommend an exhaustive exploration of the factors associated with health seeking behaviour in Ethiopia.

## Conclusion

In conclusion, our findings agreed with the findings of previous studies. The overall health seeking behavior of households was low. Specially, taking health oriented leisure activities and screening for general health was incredibly low in the community. This cues to work on promotion of healthcare on the health seeking behaviour of the population of the country. Majority of the population take action when get ill and underestimated the value of screening for general health and health oriented leisure activities. Further consideration should also be given for the risk factors including sex, marital status and level of education. Health literacy packages considering the identified differences may be designed to enhance awareness of the community about the need of health seeking behavior.

## Acknowledgements

The authors would like to thank Hosanna College of Health Sciences Research and community service for financing and supervising the overall activities of the research. We are also grateful for Hosanna town residents, data collectors and Hosanna town health office for their cooperation during the entire process of data collection.

## Supporting information

S1 Fig. Data collection tool

